# Association of pre-pregnancy body mass index with future offspring metabolic profile: findings from three independent European birth cohorts

**DOI:** 10.1101/114413

**Authors:** Diana L. Santos Ferreira, Dylan M. Williams, Antti J. Kangas, Pasi Soininen, Mika Ala-Korpela, George Davey Smith, Marjo-Riitta Jarvelin, Debbie A. Lawlor

## Abstract

**Background:** A high proportion of women start pregnancy overweight/obese. According to the developmental overnutrition hypothesis, this could lead offspring to have metabolic disruption throughout their lives, and, thus perpetuate the obesity epidemic across generations. Concerns about this hypothesis are influencing antenatal care. However, it is unknown whether maternal pregnancy adiposity is associated with long-term risk of adverse metabolic profiles in offspring, and if so, whether this association is causal, via intrauterine mechanisms, or explained by shared familial (genetic, lifestyle, socioeconomic) characteristics. We aimed to determine if associations between maternal body mass index (BMI) with offspring systemic metabolite profile are causal via intrauterine mechanisms or familial factors.

**Methods and Findings:** We used one and two-stage individual participant data (IPD) metaanalysis, and a negative-control (paternal BMI) to examine the association between maternal prepregnancy BMI and offspring serum metabolome from three European birth cohorts (offspring age at metabolite assessment 16, 17 and 31 years). Circulating metabolites were quantified by high-throughput nuclear magnetic resonance metabolomics. Results from one-stage IPD meta-analysis (*N*=5327 to 5377 mother-father-offspring trios) showed that increasing maternal and paternal BMI was associated with an adverse cardio-metabolic profile in offspring. We observed strong positive associations with VLDL-lipoproteins, VLDL-C, VLDL-triglycerides, VLDL-diameter, branched/aromatic amino acids, glycoprotein acetyls, and triglycerides, and strong negative associations with HDL-lipoprotein, HDL-diameter, HDL-C, HDL_2_-C and HDL_3_-C (all P<0.003). Stronger magnitudes of associations were present for maternal compared with paternal BMI across these associations, however there was no strong statistical evidence for heterogeneity between them (all bootstrap P >0.003, equivalent to 0.05 after accounting for multiple testing). Results were similar in each individual cohort, and in the two-stage analysis. Offspring BMI showed similar patterns of crosssectional association with metabolic profiles as for parental pre-pregnancy BMI associations, but with greater magnitudes. Adjustment of the parent BMI-offspring metabolite associations for offspring BMI suggested the parental associations were largely due to the association of parental BMI measures with offspring BMI.

**Conclusion:** Our findings suggest that maternal BMI-offspring metabolome associations are likely to be largely due to shared genetic or familial lifestyle confounding, rather than intrauterine programming mechanisms. They do not support the introduction of measures to reduce maternal BMI in order to prevent adverse offspring cardio-metabolic health.

## Introduction

In Western populations, the proportion of women who start pregnancy overweight/obese (BMI ≥25Kg/m^2^) has increased over the last 20-30 years and is now estimated to be between 20-50% [1, 2]. The developmental origin of adult diseases hypothesis proposes that maternal greater adiposity in pregnancy can prime changes in fetal metabolism that result in a life-long risk of greater adiposity and metabolic dysregulation [3]. As more adipose women have higher circulating gestational glucose, lipids and fatty acids, the fetus is purportedly *overfed,* which may lead to changes in energy metabolism and the fetal endocrine system, potentially resulting in differences in appetite control, risk of obesity, and adverse metabolism throughout the lives of offspring. This may perpetuate obesity and adverse cardio-metabolic outcomes across generations, as the daughters of overweight women would be predisposed to enter pregnancy overweight and with adverse metabolic profiles themselves. Concerns about this hypothesis are influencing antenatal care, for example recommendations related to gestational weight gain and the new criteria for diagnosing gestational diabetes are aimed at reducing future offspring obesity and adverse metabolism [4, 5]. However, whether the associations of maternal adiposity and associated traits with offspring outcomes are causal is unknown, and if they are causal, then the mechanisms are unclear [4, 6, 7].

A recent study using genetic instrumental variables (Mendelian randomization) found that intrauterine exposure to greater maternal adiposity and fasting glucose result in greater birthweight and ponderal index [6], however Mendelian randomization, within siblings analyses and negative control studies, do not support a causal intrauterine effect of greater maternal gestational adiposity on later offspring adiposity levels [8, 9]. Exposure to greater maternal adiposity and associated metabolic disruption, could nonetheless, result in more adverse metabolic profiles in offspring via intrauterine mechanisms even in the absence of an effect on offspring adiposity, for example, through a direct effect on appetite control of the development of the liver and pancreas.

The aim of this study was to examine associations between maternal pre-pregnancy BMI and multiple offspring serum metabolites in adolescence and adulthood (i.e. when offspring are in their reproductive years), using paternal BMI as a negative control. If maternal associations represent causal intrauterine effects, as opposed to being due to shared familial (lifestyle, socioeconomic, or genetic) factors linking maternal and offspring BMI and hence their metabolism, maternal associations will be stronger than paternal associations [4].

## Methods

### Study populations

All study participants provided written informed consent, and study protocols were approved by the relevant local ethics committees. The parent specific BMI associations with offspring metabolic profiles were examined in one British and two Finnish birth cohorts with nuclear magnetic resonance (NMR)-based serum metabolomics: Avon Longitudinal Study of Parents and Children [10, 11] (ALSPAC; offspring follow-up at 17 years), Northern Finland Birth Cohort 1966 and 1986 studies [12, 13] (NFBC66 and NFBC86; offspring follow-up at 31 and 16 years, respectively). Full details of the cohorts are provided in the Supporting Information Text S1 and Figure S1.

### Parental exposures and covariables

In ALSPAC: parental pre-pregnancy weight, height, education, occupation and smoking behaviour, and maternal parity were obtained during pregnancy via questionnaires. Offspring sex was obtained from obstetric records and parental and offspring ages were calculated from their dates of birth and dates of questionnaires or clinic assessments. Parental occupation was classified into social class groups from I (managerial) to IV (unskilled manual workers). Highest educational qualification for both parents was collapsed into one of five categories from none/Certificate of Secondary Education (CSE; national school exams at age 16) to university degree.

In NFBC86: parental height, weight, occupation, smoking status, offspring sex and maternal parity were collected using questionnaires given to all mothers at their first antenatal clinic visit. Level of education was obtained from questionnaires in 2001-02. Parental and offspring age was derived from their date of birth and date of assessments. Parental education was categorized into 8 categories from no occupational education to University degree, and occupation into 6 categories from entrepreneur to no-occupation.

In NFBC66: maternal height, weight, occupation, smoking status, parity, child sex were reported by mothers at the first antenatal clinic visit (16^th^ week of gestation), or in questionnaires administered between the 24^th^ and 28^th^ week of gestation. Offspring age at serum collection was derived from their date of birth and date of attendance at the 1997-1998 follow-up clinic. Maternal age in pregnancy was derived from year of birth and the date of pregnancy questionnaire completion. Education was categorized into 9 categories from none or circulating school to beyond matriculation exam and occupation into 5 categories ranging from I (highest social class) to V (no-occupation). Information on paternal BMI was not collected in NFBC66, therefore this cohort’s data was used to test for replication of the maternal-offspring association results only.

In all cohorts, head of household social class was defined as the highest occupation of either parent. For the one-stage individual participant data [14-16] (IPD) meta-analysis, education and head of house hold social class occupation categories were harmonized between cohorts (see Supporting Information Text S1 and Table S1). Individual cohort variables were used in the two-stage IPD metaanalysis. In ALSPAC, parents-offspring trios where the mother had reported that her partner was not the biological father of the offspring and those for whom this information was missing were excluded; this information was not available for the NFBCs. Differences in fetal growth in multiple pregnancies suggest that intrauterine effects are different for singletons and multiple births [17]: for the purpose of this study we considered only singleton pregnancies. For our analyses, we used data from 5327 to 5377 mother-father-offspring trios from ALSPAC and NFBC86, and 4841 to 4874 mother-offspring pairs from NFBC66 who had data on parental BMI, offspring metabolite and covariables.

### Outcomes: metabolic profiling

A comprehensive profiling of offspring circulating lipids and metabolites was done by a high-throughput NMR metabolomics platform, providing a snapshot of offspring serum metabolome at follow-up [18, 19]. In ALSPAC, offspring metabolite data were assessed on fasting (minimum 6-hours) plasma at two ages (mean age 15.5 and 17.8) and we used data from either of these. As we were interesting in lasting effects into reproductive years we prioritised measures from the older age follow-up and used the earlier measures only for participants who did not have measures at 17.8. Mean age at assessment in the whole cohort after using both time points was 17. Participants of both NFBCs fasted overnight before serum collection on the morning of clinic attendance (8 to 11am) at mean age 16 (NFBC86) and 31 years (NFBC66).

Collectively, the 153 metabolites measured by the platform represent a broad molecular signature of systemic metabolism [18, 19]. The platform provided simultaneous quantification of lipoprotein lipids and subclasses, fatty acids and fatty acid compositions, ketone bodies, amino acids, as well as glycolysis and gluconeogenesis-related metabolites in absolute concentration units. This platform has been applied in various largescale epidemiological and genetic studies [20-23]; the detailed protocol, including information on quality control, has been published elsewhere [19, 24] and more information is given in the Supporting Information.

### Statistical analysis

Data cleaning and checking were performed separately for each cohort. One-stage and two-stage IPD [14-16] meta-analyses were performed to assess the associations of maternal pre-pregnancy BMI with offspring metabolic profiles, using paternal BMI as a negative control. The term *IPD* relates to the data recorded for each participant in a study [14]. Linear regression models were adjusted for parental age, smoking, education, head of household social class, maternal parity, offspring age at blood collection and sex. Robust standard errors were estimated for all associations and probability values as some metabolite concentrations had skewed distributions.

We conducted three sets of IPD meta-analysis that included maternal vs paternal comparisons:

1. A one-stage IPD meta-analysis restricted to the two cohorts with data on both maternal and paternal BMI (ALSPAC and NFBC86) was considered to be our main analyses. In these analyses offspring metabolites were standardized (z-scored) across both cohorts and then regressed on maternal and paternal z-scored BMI (again standardized across both cohorts) with adjustment for the harmonized covariables and a binary variable reflecting whether the participant was from ALSPAC or NFBC86. These analyses were conducted on between 5327 and 5377 trios (numbers with complete data varied slightly for different metabolites). This approach has greater statistical efficiency than a two-stage IPD meta-analysis, and restricting to the two cohorts with both maternal and paternal BMI ensures that exposure-outcome comparisons are possible within all parental-offspring trio data available. It assumes that the two cohorts are from the same population to which inferences are being made.
2. A one-stage IPD meta-analysis, undertaken as described above, but including the maternal BMI-offspring metabolite associations from NFBC66. This sensitivity analysis assessed associations of maternal BMI with offspring metabolites in up to 4874 additional mother-offspring pairs (10,181 to 10,251 in total) from ALSPAC, NFBC86 and NFBC66 with the same associations of paternal BMI with offspring metabolites in ALSPAC and NFBC86 only (N = 5327 to 5277). This has greater statistical power for the maternal associations and for determining differences between mothers and fathers. It assumes that all three cohorts are from the same population to which inference is being made. This includes the assumption that paternal BMI, would be similarly associated with offspring metabolites in NFBC66 as in the other two cohorts, if we would have been able to model data on this unmeasured trait.
3. Two-stage IPD meta-analysis, in which the regression outputs of standardized offspring metabolites with standardized parental BMI were derived separately for each cohort and for maternal and paternal BMI (stage 1), and then pooled results from each cohort were meta-analysed together using the random effects inverse-variance-weighted method (stage 2). The standardization of offspring metabolites and parental BMI was undertaken within each cohort in this method. We used this method to: (a) explore whether our standardization across cohorts of offspring metabolites and parental BMI and harmonization of potential confounders in the one-stage IPD meta-analyses had notably influenced results; (b) to test for heterogeneity in association results between three cohorts (two for paternal BMI-offspring metabolite associations) using the I^2^ statistics [25], which provide a test of replication of our findings and tests our assumption that the cohorts are likely to be from the same population to which we want to make inference and (c) by using a random effects method for pooling results (even if there was little evidence of heterogeneity) we tested whether results were similar if we relaxed the assumption of the cohorts all being from the same underlying population (this approach provides results that are interpreted as the average across studies assuming that these might reflect different populations).

Magnitudes of maternal and paternal associations were compared by linear fit. In addition, the magnitudes of one-stage IPD associations were compared to the corresponding two-stage IPD for maternal and paternal BMI separately, also by examining their linear fit. The differences between maternal and paternal associations (in all three approaches) were calculated from the bootstrap replicate distribution (1000 replications). Beta-estimates and standard errors were empirically calculated from the mean and SD of the bootstrap distribution, respectively. All p-values were calculated using bootstrap means and standard errors and compared to a z-distribution.

To establish a threshold that takes into account multiple testing and the correlation structure of the metabolite data, principal components analysis (PCA) was carried out on the z-scored metabolite data [26]. The first 17 principle components (PCs) explained 95% of the variance, this number is a proxy of the number of independent tests being performed. Therefore a threshold was defined as p-value<0.003 (i.e. α/17 where α=0.05) for assessing associations with the 153 metabolites.

We performed two additional analyses using the one-stage IPD meta-analysis only that aimed to explore whether any maternal (or paternal) pre-pregnancy BMI associations with offspring metabolites were mediated by the relationship of the parent’s BMI with offspring BMI. First, we examined the associations of offspring BMI (assessed at the same time as blood collection) with their metabolites. Second, we repeated the meta-analysis described above, in point 1, with additional adjustment for offspring BMI. Both analysis were undertaken in N=5266 to 5316 trios from ALSPAC and NFBC86 cohorts.

Statistical analyses were conducted using R version 3.0.1 (R Foundation for Statistical Computing, Vienna, Austria) and Stata version 14.1 (Stata Inc., TX, USA).

## Results

Characteristics of the three study populations are shown in Table S1 and the flowchart in Figure S1. The percentage of overweight/obese mothers was 20%, 15% and 23% in ALSPAC, NFBC86 and NFBC66, respectively and of fathers was 47% and 32% in ALSPAC and NFBC86, respectively. The highest percentage of overweight/obesity in offspring at follow-up in adolescent or adulthood was seen in NFBC66 (mean age 31 y) at 40%, followed by ALSPAC with 21% (mean age 17 y), and NFBC86 with 11% (mean age 16 y).

Figure 1 shows associations of parental BMI with the 153 offspring metabolic measures, each expressed as a difference in means in SD units for a 1-SD greater parental BMI, from a one-stage IPD meta-analysis restricted to the two cohorts with complete trio data (N= 5327 to 5377 trios). Table S2 shows the same associations expressed as magnitudes in absolute concentration units (e.g. mmol/l per 1-SD difference in parental BMI). Both maternal and paternal pre-/early-pregnancy BMI were associated with a large proportion of the offspring metabolites (maternal BMI associated with 49% and paternal BMI with 44% of offspring metabolites at p<0.003). The associations were mostly in the same direction for both parents and were in the direction of more adverse cardio-metabolic risk in offspring with higher parental BMI in pregnancy.

**Figure 1.**
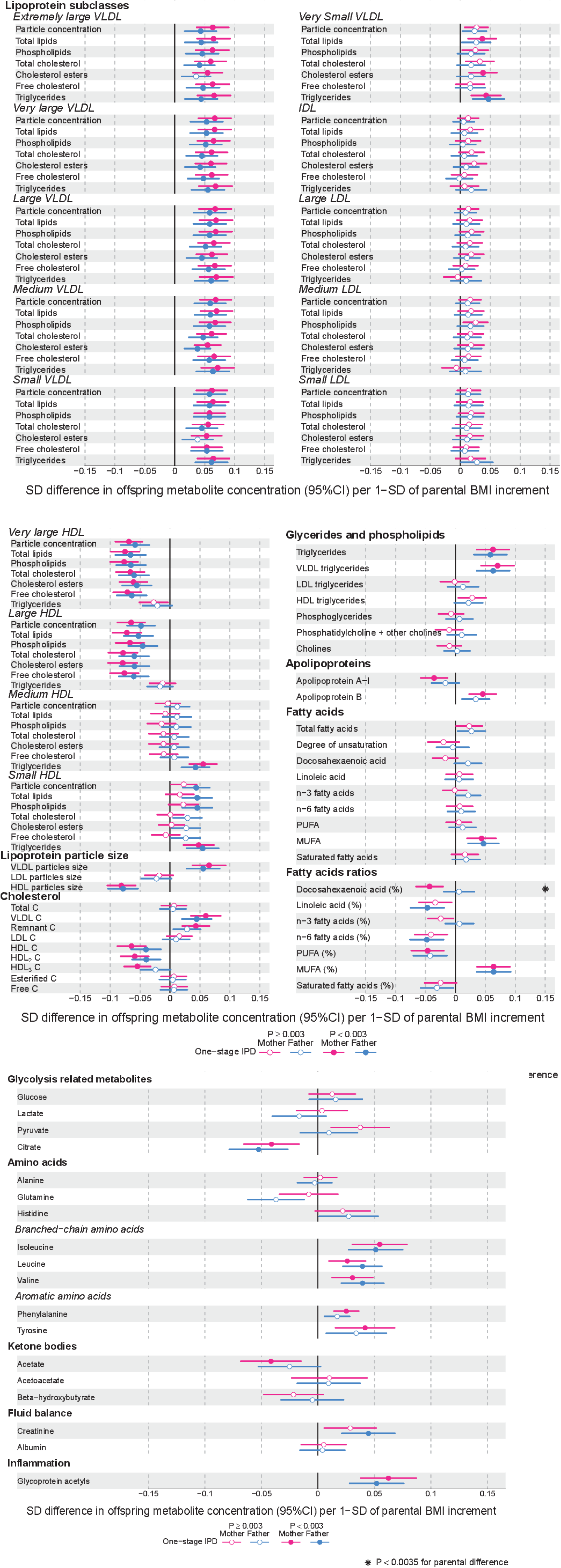
One-stage individual participant data meta-analysis: offspring lipoprotein, lipids and metabolite differences in means in SD units per 1-SD higher maternal (pink) or paternal (blue) BMI, meta-analysed across ALSPAC and NFBC86 cohorts.

*Associations were adjusted for parental age, smoking, education, head of household social class, maternal parity, offspring’s age at blood collection, sex and cohort membership. Results are shown in SD-scaled concentration units of outcome, differences in absolute concentration units are listed in Table S2. Error bars= 95% confidence intervals (CI). VLDL=very-low-density lipoprotein; IDL=intermediate-density lipoprotein; LDL=low-density lipoprotein; HDL= high-density lipoprotein; C= cholesterol; MUFA=monounsaturated fatty acids; PUFA=polyunsaturated fatty acids.*

Amongst lipoprotein lipids, the most marked positive associations were observed in five VLDL subclasses (all VLDL except the *very small* sub-class). There were no robust associations of parental BMI with offspring IDL and LDL subclasses. Strong inverse associations were observed for very large and large HDL subclasses with the exception of structural triglycerides that exhibited no clear associations. Lipoprotein particle sizes followed the same pattern of associations as those observed with lipoprotein subclasses. Parental BMI was strongly positively associated with offspring VLDL cholesterol, VLDL triglycerides, triglycerides and remnant cholesterol, with the same direction and comparable magnitudes to the first five VLDL lipoprotein subclasses. Conversely, HDL, HDL2, and HDL3 cholesterol sub-fractions showed inverse associations, with magnitudes similar to very large and large HDL lipoprotein subclasses. Associations of parental BMI with apolipoprotein A-I and B were in opposite directions, in agreement with the association pattern for lipoprotein subclasses.

Parental BMI was strongly positively associated with offspring branched-chain amino acids, and maternal BMI with aromatic amino acids.

Mono-unsaturated fatty acids (MUFA) was the only fatty acid clearly associated with parental BMI when assessed alone or as a proportion of total FAs (MUFA-to-total FA ratio). Several of the other FAs showed associations with parental BMI, but only as proportions of total FAs. Poly-unsaturated (PUFA) to total fatty acids (PUFA-to-total FA ratio), ratio of linoleic acid (LA) to total fatty acids (LA-to-total FA ratio) and omega 6 to total fatty acids ratio were all inversely associated with parental BMI.

Parental BMI was not clearly associated with any of the glycolysis related metabolites, except citrate were there was an inverse association. Parental BMI was strongly positively associated with offspring glycoprotein acetyls, an inflammatory marker.

Overall, associations across offspring metabolites were similar for maternal and paternal BMI entered as exposures (Figure 2; R^2^= 0.89 and slope= 0.78 ± 0.02): for the majority there was no strong statistical evidence that associations differed other than what might be expected by chance (*P*_boot_ >0.003). DHA-to-total FA ratio was the only metabolite that showed some suggestion of difference in association between maternal and paternal BMI (mean difference in % per 1-SD BMI: β_mother_=-0.01, 95% CI_mother_=−0.02 to −0.01, *P*_mother_=0.0002 *versus* β_father_=0.002, 95% CI_father_= −0.01 to 0.01, *P*_father_=0.66; *P*_boot_ for difference between the two associations = 0.0034; Figure 1). However, the positive maternal association and the difference between maternal and paternal association appeared to be driven primarily by results in ALSPAC as the difference was not seen in NFBC86 or the two-stage IPD metaanalysis (Figure S2).

**Figure 2.**
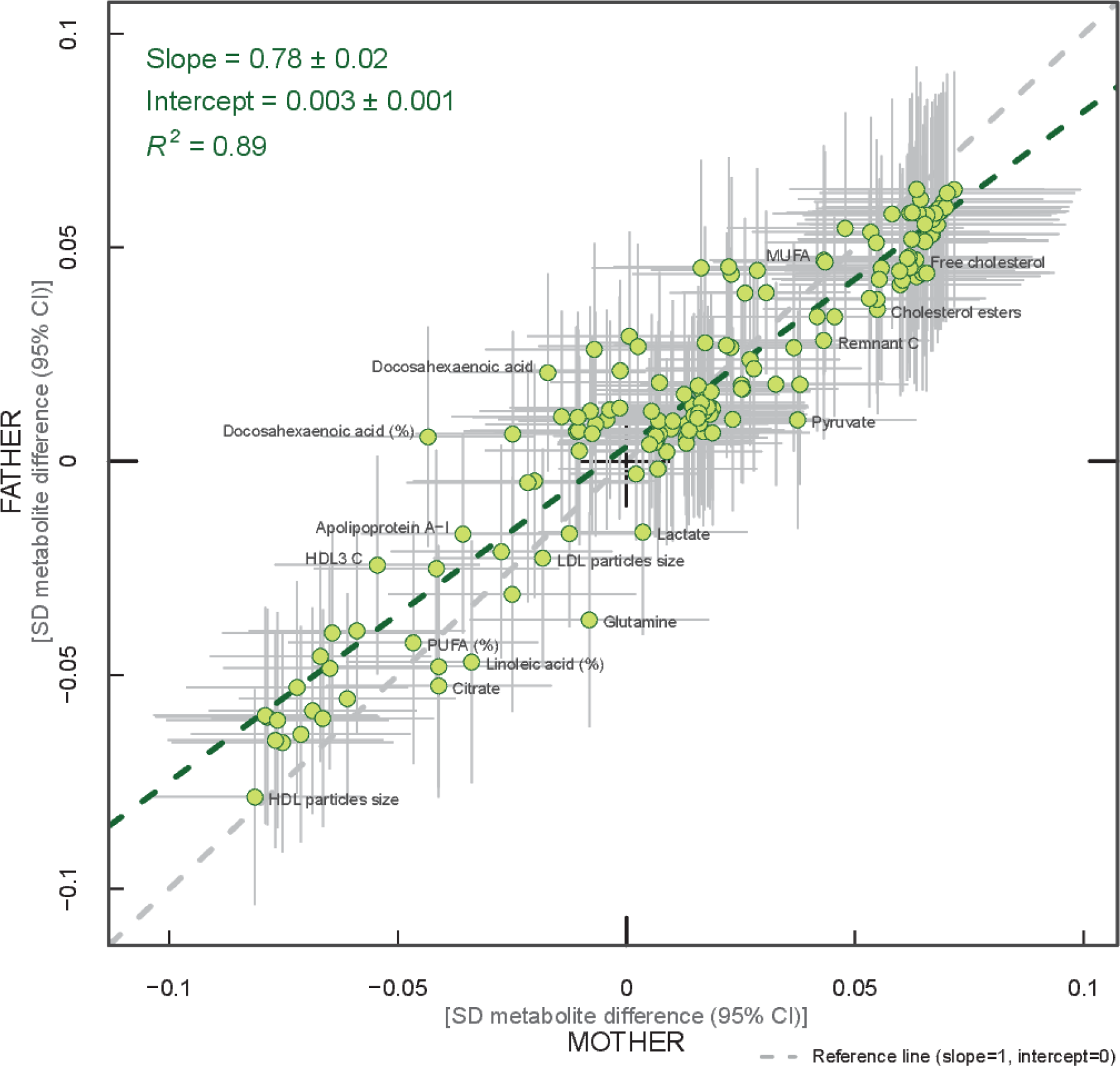
Linear fit between paternal and maternal models (green dashed line). R^2^ indicates goodness of fit.

*Each green dot represents a metabolite and the positions of the dots are determined by difference in mean offspring metabolite (in SD units) for each increase of 1SD maternal BMI (x-axis) and difference in mean offspring metabolite (in SD units) for each increase in 1SD paternal BMI (y-axis). The horizontal grey lines on each dot denote the confidence intervals (CI) for maternal associations and the vertical grey lines indicate the CI for paternal estimates. A linear fit of the overall correspondence summarizes the similarity in magnitude between maternal and paternal associations (green dashed line). A slope of 1 with an intercept of 0 (dashed grey line), with all green dots sitting on that line (R^2^=1), would indicate that maternal and paternal estimates had the same magnitude and direction. Results are shown in SD-scaled concentration units of outcome, difference in absolute concentration units are listed in Table S2.*

When the one-stage analyses were repeated to compare maternal BMI-offspring metabolite associations in all three cohorts (*N*= 10,181 to 10,251) to paternal BMI-offspring metabolite associations in the two cohorts with paternal BMI (*N*= 5327 to 5377), the maternal associations were essentially the same as those described above; no statistical evidence for stronger maternal associations emerged (Figure S3).

In the two-stage meta-analysis, the associations were highly consistent across the three cohorts, with just 5% and 3% of associations having I^2^ statistic ≥75% for maternal and paternal BMI, respectively (Figure S2 and >Table S3). A comparison of the results from one and two-stage IPD meta-analyses showed very similar patterns of association for both maternal and paternal BMI (R^2^= 0.96 and slope=0.81 ± 0.01; R^2^=0.99 and slope=1 ± 0.01, respectively for maternal and paternal BMI; Figure S4). The magnitudes of associations in absolute concentration units for individual cohorts, as well as for the two-stage IPD meta-analysis are shown in Tables S3-S5.

Cross-sectional associations of offspring BMI with their metabolite concentrations had very similar patterns (in terms of direction and which metabolites had strongest and weakest associations) to those seen for parental BMI, but were considerably stronger in magnitude (Figure S5). When adjusting parental BMI-offspring metabolite associations for offspring BMI, most attenuated markedly, such that many point estimates were in the opposite directions and the vast majority were consistent with the null following this adjustment (Figure S6).

## Discussion

We report the first study that investigates the potential influence of maternal pre-pregnancy BMI on adult offspring serum metabolome, in order to determine whether intrauterine mechanisms, related to developmental overnutrition, result in metabolic disruption in adults when they are in their reproductive years. We found similar associations of both maternal and paternal BMI with offspring systemic metabolism 16-17 years later and 31 years later (the latter assessed with maternal BMI only), suggesting that shared familial genetic, socioeconomic and lifestyle characteristics, rather than an intrauterine programming effect explain the associations of maternal pre-pregnancy BMI with offspring metabolites.

Associations of greater parental BMI with offspring lipids were in the directions of an adverse cardio-metabolic profile, with positive associations with VLDL-lipoproteins, VLDL-C, VLDL-triglycerides, VLDL-diameter, branched/aromatic amino acids, glycoprotein acetyls, and triglycerides, and inverse associations with HDL-lipoprotein, HDL-diameter, HDL-C, HDL_2_-C and HDL_3_-C. Associations were also seen with FA ratios and some amino acids. In further analyses we demonstrated that in addition to similar associations between paternal BMI with offspring metabolites (to those seen for maternal BMI) the offspring’s own BMI was associated in a similar pattern (but with stronger magnitudes) to their metabolites, further supporting the notion that shared familial characteristics explain these associations. Lastly, we found that parental associations with metabolites attenuated markedly with adjustment for offspring BMI, suggesting that shared familial characteristics drive associations of maternal pre-pregnancy BMI and paternal BMI with their offspring BMI, which in turn results in metabolic disruption. However, we acknowledge that we cannot rule out the possibility of a small intrauterine causal effect of higher maternal adiposity during pregnancy on offspring metabolism, which we may have been unable to detect despite our large sample size.

Previous studies have compared associations of maternal and/or paternal BMI, measured pre/early-pregnancy, with offspring adiposity (BMI [3, 27-42], overall adiposity [27, 32, 36, 38, 42, 43], central adiposity [27, 28, 32, 36, 38, 44], lean mass [32, 43]), with fewer also examining association with offspring lipids (HDL-C [27, 28, 42, 44, 45], LDL-C [27, 42], total cholesterol [27, 28, 42, 45],triglycerides [27, 36, 44]), blood pressure [27, 28, 36, 44], glucose [44] and insulin [27, 36, 42]. Our study is generally larger than these previous studies, examines a detailed metabolic profile with 153 metabolite concentrations and with outcomes assessed in offspring at older ages. Where it is possible to make comparisons, our findings are consistent with the reported lack of association between maternal BMI and offspring total cholesterol [27, 28, 42, 45], LDL-C [27, 42] and glucose [44] and inverse association with HDL-C [27, 44] (previous studies *N=70* to 4871; offspring age range: 4-8 y). Our results also contrast reports from some of the same studies that found no association of maternal BMI with offspring triglycerides [27, 36, 44] or HDL-C [28, 42, 45]. These differences might reflect differences in the age of offspring outcome and/or that we are able to assess associations with more refined outcomes (e.g. HDL-C sub-fractions). The only previous study that we were able to identify that compared maternal BMI- to paternal BMI- associations with any outcomes similar to ours and with a similar size (N=4871) to our main analysis approach, reported similar associations of maternal BMI with total cholesterol, LDL-C and HDL-C to those seen for paternal BMI [27], as in our study, though that study found no association of either parents BMI with offspring triglycerides. Association between offspring BMI and their serum metabolomics profile are similar to the ones reported by Wurtz et al. [46].

Our study has several strengths. It has a large sample size, included replication testing across three different birth-cohorts from two different countries and very detailed offspring blood metabolome measurements. The consistency of associations across three independent studies and using different analytical approaches suggests that our results are unlikely to be due to chance. Furthermore, we used a negative-control approach (with paternal BMI) to explore causal inference. Using paternal BMI as a negative-control makes it possible to disentangle the extent to which maternal BMI-offspring metabolite associations are due to casual intrauterine effects or confounding by familial factors (i.e. genetic and/or shared lifestyle traits within families) [4, 47]. We performed a one-stage IPD metaanalysis, which is the *gold standard* of meta-analysis [48], for our main analytical approach, but also tested the assumptions of this approach, and the effect of harmonisation across studies, by comparing the results to a two-stage IPD, and found high levels of consistency between the two approaches.

Limitations of our study include the use of parental BMI as a measure of adiposity; different body composition characteristics, such as fat distribution or fat to lean body mass ratio, may show differential associations with offspring serum metabolome. Moreover maternal BMI was selfreported, and paternal BMI was reported by mothers, which might have led to misclassification of parental BMI. Our findings cannot be used to draw inferences about the role of maternal specific nutritional intakes, such as glucose, lipids and free fatty acids, during intrauterine period on later offspring metabolic profile, as these were not measured in our cohorts during pregnancy. It is possible that non-paternity for some of the fathers has affected our results (for the NFBCs we had no information on this possibility). However, any impact of non-paternity would be likely to selectively reduce paternal BMI associations, since there would be no shared genetic effects on the association of paternal BMI with offspring metabolites, and family lifestyle and socioeconomic associations may be weaker between non-biological parents and offspring. This would thus tend to enhance maternal-paternal differences, whereas we see similar associations between parents. Despite adjusting for several potential confounders, residual confounding may still explain the associations that we have observed between parental BMI and offspring. For example, we were unable to adjust for parental physical activity or dietary intake. These confounders are likely to be similar for maternal and paternal BMI and so would unlikely bias our inference about specific maternal effects on offspring metabolism (potentially due to intrauterine programming). Whilst we have a large sample size, we may lack power to detect small differences between maternal and paternal BMI associations with offspring metabolites, especially as the human circulating metabolome is a tightly controlled homeostatic system, and even small differences between some metabolites might have important clinical impact. We have assessed BMI as a continuous variable to explore the hypothesis that each incrementally greater maternal BMI overfeeds their developing infant in utero in a dose-response way and our results suggest that if that is the case it has no long-term effect on offspring metabolism. However, we cannot exclude a threshold effect – i.e. maternal obesity (or extreme obesity) having a long-term effect via intrauterine mechanisms. Furthermore, our results may not necessarily generalise to other non-European populations.

In conclusion, the similarity of association between pre-pregnancy maternal BMI and paternal BMI with offspring metabolic profiles, suggest that maternal BMI associations are likely to be largely due to shared genetic or familial lifestyle confounding, rather than intrauterine mechanisms. These findings do not support the introduction of antenatal measures to reduce maternal pregnancy BMI in order to prevent disruption of offspring cardio-metabolism in later life. Interventions to reduce BMI in all family members may be more beneficial.

## Abbreviations

ALSPAC: Avon Longitudinal Study of Parents and Children;
BMI: body mass index;
C: cholesterol;
FA: fatty acids;
HDL: high-density lipoprotein;
IDL: intermediate-density lipoprotein;
IPD: individual participant data;
LA: linoleic acid;
LDL: low-density lipoprotein;
MUFA: mono-unsaturated fatty acids;
NFBC66: Northern Finland Birth Cohort of 1966;
NFBC86: Northern Finland Birth Cohort of 1986;
NMR: nuclear magnetic resonance;
PCA: principal component analysis;
PUFA: polyunsaturated fatty acids;
SD: standard deviation;
VLDL: very-low-density lipoprotein.

## Acknowledgments

ALSPAC: We are extremely grateful to all the families who took part in this study, the midwives for their help in recruiting them, and the whole ALSPAC team, which includes interviewers, computer and laboratory technicians, clerical workers, research scientists, volunteers, managers, receptionists and nurses.

NFBC66, NFBC86: The authors are very grateful to all the participants who took part in the Northern Finland Birth Cohort 1966 and 1986 study, to the whole study team, including research staff and all others involved in data collection and processing, and to those in the oversight and management of the study.

## Supporting Information

***Figure S1.*** Overview of the study design, cohorts and statistical analyses. IPD= individual participant data; analysis 1 is our main analysis.

(PDF)

***Figure s2.*** *Two-stage IPD meta-analysis and individual cohort associations: offspring lipoprotein, lipids and metabolite differences per 1-SD higher maternal (pink) or paternal (blue) BMI.*

(PDF)

***Figure S3.*** *One-stage IPD meta-analysis: offspring lipoprotein, lipids and metabolite differences in means in SD units per 1-SD higher maternal (pink) or paternal (blue) BMI, meta-analysed across ALSPAC, NFBC86 and NFBC66 cohorts.*

(PDF)

***Figure S4.*** *Linear fit between two and one-stage individual participant data (IPD) metaanalysis for mother (left panel; pink dashed line) and father (right panel; blue dashed line) models.*

(PDF)

***Figure S5.*** One-stage IPD meta-analysis: offspring lipoprotein, lipids and metabolite differences in means in SD units per 1-SD higher maternal (pink), paternal (blue) or offspring (green) BMI, meta-analysed across ALSPAC and NFBC86 cohorts.

(PDF)

***Figure S6.*** One-stage IPD meta-analysis: offspring lipoprotein, lipids and metabolite differences in means in SD units per 1-SD higher maternal (pink) or paternal (blue) BMI, meta-analysed across ALSPAC and NFBC86 cohorts, with and without further adjustment for offspring BMI.

(PDF)

***Table S1.*** Characteristics of the three study populations.

(PDF)

***Table S2.*** One-stage individual participant data (IPD) meta-analysis: offspring lipoprotein, lipid and metabolite absolute concentration differences per 1-SD higher parental BMI.

(PDF)

***Table S3.*** Two-stage IPD meta-analysis: offspring lipoprotein, lipid and metabolite absolute concentration differences per 1-SD higher parental BMI.

(PDF)

***Table S4.*** *ALSPAC: offspring lipoprotein, lipid and metabolite absolute concentration differences per 1-SD higher parental BMI.*

(PDF)

***Table S5.*** *NFBC66 and NFBC86: offspring lipoprotein, lipid and metabolite absolute concentration differences per 1-SD higher parental BMI.*

*(PDF)*

***Text S1.*** *Supplemental methods.*

(PDF)

